# Fitness effects and transmission of phoretic nematodes of the burying beetle, *Nicrophorus vespilloides*

**DOI:** 10.1101/193714

**Authors:** Yin Wang, Daniel E. Rozen

## Abstract

1. *Nicrophorus vespilloides* is a cosmopolitan social beetle that rears its offspring on decomposing carrion. Wild beetles are frequently associated with two types of macrobial symbionts, mites and nematodes.
2. Although these organisms are believed to be phoretic commensals that harmlessly use beetles as a means of transfer between carcasses, the role of these symbionts on *N. vespilloides* fitness is poorly understood. Here we show that nematodes have significant negative effects on beetle fitness across a range of worm densities and also quantify the density-dependent transmission of worms between mating individuals and from parents to offspring.
3. Using field-caught beetles, we provide the first report of a new nematode symbiont in *N. vespilloides*, most closely related to *Rhabditoides regina*, and show that worm densities are highly variable across individuals isolated from nature but do not differ between males and females. Next, by inoculating mating females with increasing densities of nematodes, we show that worm infections significantly reduce brood size, larval survival and larval mass, and also eliminate the trade-off between brood size and larval mass. Finally, we show that nematodes are efficiently transmitted between mating individuals and from mothers to larvae, directly and indirectly via the carcass, and that worms persist through pupation.
4. These results show that the phoretic nematode *R. regina* can be highly parasitic to burying beetles but can nevertheless persist because of efficient mechanisms of intersexual and intergenerational transmission.
5. Phoretic species are exceptionally common and may cause significant harm to their hosts, even though they rely on these larger species for transmission to new resources. However, this harm may be inevitable and unavoidable if transmission of phoretic symbionts requires nematode proliferation. It will be important to determine the generality of our result for other phoretic associates of animals.

## Introduction

Animals that persist on ephemeral and spatially dispersed resources have evolved diverse mechanisms to detect and exploit these resources (1–3). Carrion feeders, like blowflies and burying beetles, can use olfactory cues to detect minute concentrations of the volatile products of animal decomposition and can orient their search flights accordingly (4, 5). However, some animals are incapable of moving across large distances themselves. Instead, these species hitch a ride on the bodies of other more mobile species and are consequently transported from resource to resource (6, 7) Thus rather than developing mechanisms to detect resources, they have evolved mechanisms to ensure reliable and durable associations with the species that carry them (8, 9). This strategy, known as phoresy, is common in many species of insects, mites and nematodes and is a form of symbiosis that is typically believed to be harmless to the host (10, 11). The rationale for this belief is that because phoretic species are wholly dependent on their hosts for their migration, species that cause too much harm and thereby reduce their transport between breeding resources, face the risk of local extinction (11). However, just as parasites and pathogens can evolve levels of virulence that balance harm to hosts with the need to be transmitted between hosts, so too may phoretic species become parasitic, as long as this harm facilitates their transmission between hosts (12–14). Few studies have quantified the direct harm of phoretic species to their hosts while also estimating their persistence and transmission between host individuals and across generations. Our aim in this paper is to address these questions in the context of the burying beetle, *Nicrophorus vespilloides*, and its phoretic nematodes.

*Nicrophorus* burying beetles are subsocial insects that breed on small vertebrate carrion (15). After locating a small vertebrate carcass using volatile cues produced from microbial decomposition (5), a mated female lays eggs in the surrounding soil after which she (or a mated pair) prepare the carcass for the arrival of the hatched larvae (16). The carcass is buried underground, stripped of fur or feathers, the gut is removed, and then it is coated in antimicrobial oral and anal secretions (17–21). When larvae migrate to the carcass, parents remain to feed them via regurgitation (22, 23), which both provides a meal and also transmits the endogenous microbiome to the developing larvae (24–27). But beetles are not alone in their consumption of the carcass. *Nicrophorus* adults trapped in the field are conspicuously associated with high densities of mites and nematodes that are attached to their carapace or reside internally (28, 29). Many species of mites have established phoretic associations with burying beetles, and their well-studied effects on beetles range from harmful to beneficial, depending on the context and the study (30, 31). By contrast, only one species of phoretic nematode has been described in *Nicrophorus* and its effects on beetles are unknown (29).

Richter (1993) described the carrion-feeding nematode *Rhabditis stammeri* isolated from *N. vespilloides*. He showed that worms were present in the gut and genitalia and could be transmitted between mating individuals (29). However, although *Nicrophorus* researchers regularly comment on the presence of nematodes in laboratory and field populations, there is no direct evidence that these worms are actually *R. stammeri*, nor is there any understanding of their natural abundance in field-caught insects. More importantly, we lack an experimental understanding of the fitness consequences of these nematodes for beetles. As part of our efforts to understand the evolution and ecology of phoretic associates of *N. vespilloides* we provide a detailed study of the identity and effects of a novel nematode associate of *N. vespilloides,* most closely related to *Rhabditoides regina*, that we cultured and quantified from field-caught beetles. In brief, we find that these nematodes are extremely numerous in wild beetles and significantly reduce *N. vespilloides* fitness. In addition, worms are efficiently transmitted in high densities between mating adults and from infected mothers to their offspring, which then persist through beetle development and are retained into adulthood. We discuss these results in the context of the evolution of interspecific interactions in *N.vespilloides* and the evolution of host harm in phoretic species.

## Methods

### General procedures

All experimental beetles were taken from an outbred laboratory population derived from wild-caught *N. vespilloides* individuals trapped in Warmond, near Leiden in The Netherlands, between May and June 2016. Beetles were maintained in the laboratory at 20°C with a 15:9 hour light:dark cycle. All adults were fed fresh chicken liver twice a week. To maintain the laboratory population and to establish experimental broods, an unrelated male and female were placed overnight without food in a small plastic container filled with 1-2 cm of autoclaved soil for mating. The following morning, mated females were provided with a freshly thawed mouse carcass (20-23g) in a new larger container for egg laying. Broods were reared until larvae dispersed from the carcass, approximately 7 days post-hatching (32). Dispersed larvae were placed together into a new container of sterile soil until eclosion, at which point they were removed to new individual containers.

To generate nematode-free adults, eggs were collected from broods within 12-24 hours of laying and surface sterilized with an antimicrobial solution of hen egg white lysozyme (1 mg/ml), streptomycin (500 μg/ml) and ampicillin (100 μg/ml) (33). These were then transferred onto 1% water agar plates to hatch. Subsequent microscopy confirmed that treated eggs are free of nematodes. To prevent nematode transmission from parents to newly hatched larvae, 1^st^ generation nematode-free larvae were reared without parental care on freshly thawed mouse carcasses. Once these nematode-free individuals had eclosed as adults, they were crossed as above, and maintained thereafter on autoclaved soil.

### Nematode quantification from field-caught and lab beetles

Nematodes were collected and counted from the guts and cuticles of field-caught and lab-reared beetles. Individual adult beetles were vortexed for 3 minutes in 700 μl sterile phosphate buffer saline (PBS, pH= 7.2) to collect nematodes from the cuticle. To quantify nematodes from the beetle gut, we removed individual beetle guts with fine forceps and suspended these in 700 μl sterile PBS. 10 μl of each suspension (cuticle or gut sample) was then transferred onto a haemocytometer and examined at 10X magnification for counting. Three independent 10 μl aliquots were counted from each sample to generate a mean value estimate/sample.

### Nematode maintenance and identification

Experimental nematodes were isolated directly from the cuticles of field-caught beetles. Species identification is explained below. To maintain laboratory populations, newly collected nematodes were transferred onto petri plates containing Nematode Growth Medium (NGM contains: 1.7% agar/L; 50mM NaCl; 0.25% peptone; 1mM CaCl_2_; 5μg/mL Cholesterol; 25mM KH_2_PO_4_; 1mM MgSO_4_) (34) and fed with an *E. coli* strain originally isolated from a mouse carcass and held at 20°C. Nematodes were transferred to fresh plates containing *E. coli* at an intial density of ∼10^6^ cells/plate every 2 days.

Nematode samples reared on NGM plates were collected and suspended in sterile PBS (100mM, pH 7.2), after which they were surface sterilized in a wash solution containing a 1:2 ratio of 5N NaOH and a 5% solution of sodium hypochlorite (34). Washed nematodes were re-suspended in 1ml PBS, and then centrifuged at 13,000 × g for 10 mins. Nematode pellets were re-suspended in 0.7 ml PBS and stored at -20 °C. For DNA extraction, nematode samples were thawed and homogenized with a sterile micropestle and vortexed for 2 mins. Samples were then lysed in SDS at 60°C for 30 min. following the method of Donn *et al*. (35). DNA was extracted using phenol-chloroform and quantified using a Thermo NanoDrop D-1000 spectrophotometer. The 18S rRNA gene fragment (∼ 900 bp) was amplified using primer pairs Nem_18S_F [Code] and Nem_18S_R[Code] (36). For PCR amplification, 2μl of template containing 2-10 ng DNA was used directly in a 20μl reaction mixture using *Pycoccus furious* (*Pfu*) DNA Polymerase. PCR was performed in a thermal cycler (Bio-RAD T100^TM^) with thermal cycling of 95 °C for 5 min, followed by 35 cycles of 95 °C for 30 sec, 54 °C for 30 sec, 72 °C for 30 sec and a final extension at 72 °C for 5 min. PCR products (fragment length of 900bp) were gel purified (illustra^TM^ GFX^TM^ PCR DNA and Gel Band Purification Kit) and sequenced commercially via MaxyGen. The resulting 18s rRNA gene sequence was classified to species using a nucleotide BLAST against the NCBI database.

### Fitness effects and transmission of nematode infections

To determine the fitness effects of nematodes on beetles, broods were established with worm-inoculated mated females (at least 20 broods/inoculation density). Before inoculation, nematodes were surface sterilized to remove any surface-associated bacteria, and then suspended in sterile PBS. Experimental beetles were inoculated with either ∼10, 10^2^, 10^3^ and 10^4^ nematodes per beetle by pipetting worm solutions under their elytra and on their natural openings (mouth, spiracles and anus). Two days later, inoculated females were paired with an unrelated worm-free male and allowed to mate overnight in small plastic containers. The next morning, mated females were provided with a freshly thawed mouse carcass and allowed to rear their broods until the point of larval dispersal. When the beetle larvae dispersed from the carcass, we measured the brood size and total brood mass for each brood.

To quantify nematode transmission between *Nicrophorus* individuals, we measured the number of worms transmitted between experimentally inoculated beetles and worm free recipients. Transmission was examined between mating adults and from mothers to offspring.

Nematode transmission was quantified bi-directionally between males and females (i.e. female donors to male recipients and male donors to female recipients). Individual adults were first inoculated with different worm densities, as outlined above, and then maintained for 2 days in small boxes containing 1-2 cm of sterile soil. Next, these individuals were transferred to a new box with sterile soil, and paired with a nematode-free individual of the opposite sex for mating. Two days later, both individuals were sampled to determine nematode densities.

To estimate worm transmission from parental females to offspring, females were inoculated with different worm densities, allowed to mate with a worm-free male, and then provided a fresh carcass for breeding. When beetle offspring eclosed, they were sampled to estimate nematode densities.

## Results

### Nematode identification and infection densities in wild beetles

The partial nematode 18S rRNA gene sequence (900bp) was BLASTed against the NCBI database and showed 95% identity to *Rhabditoides regina* strain DF5012 (AF082997) and was clearly distinct from *Rhabditis stammeri*, the nematode species already described from *N. vespilloides*. With the resolution we have from this sequence, it remains uncertain if our isolate is truly *R. regina* or an as yet undescribed species; further sequencing will be required in a later study to more fully resolve its taxonomy. For ease of presentation we hereafter tentatively refer to our isolate as *R. regina.* Although *R. regina* has not been previously reported in *Nicrophorus*, it has been reported as a parasite in scarabaeid beetle larvae (37).

Nematode densities were quantified from field-caught beetles, and we observed no overall differences in densities between males and females or between the gut and cuticle samples (all tests NS) (Figure 1). The mean number of nematodes in females was (Mean ± SE: 1,720.47 ± 828.45) and for males was (Mean ± SE: 978.27 ± 372.75). The number of nematodes in females and males was highly variable (females: 40 - 12,101; males: 10 - 4,608 worms).

**Figure 1.**
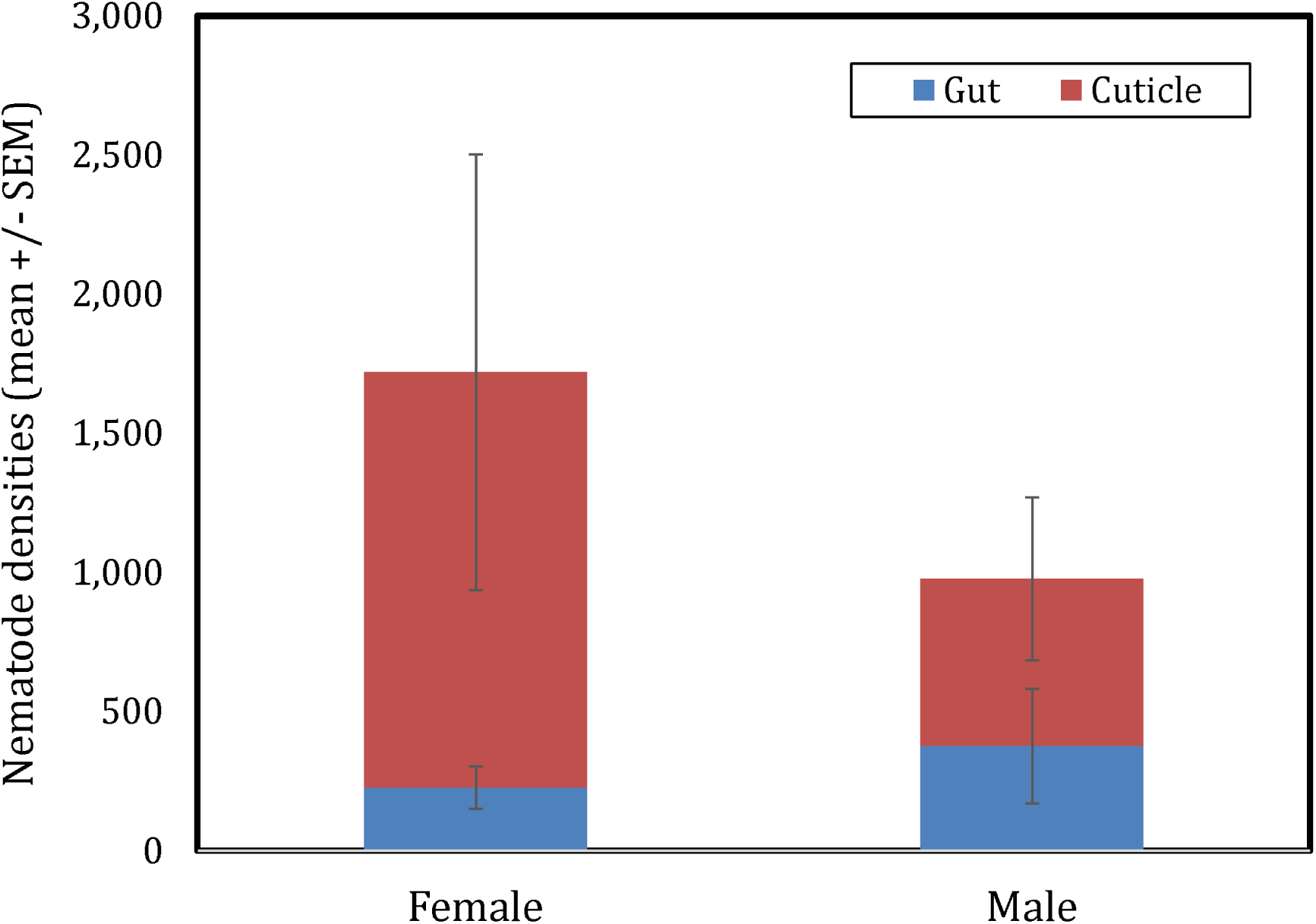
Nematode densities in field-caught *N. vespilloides.*

### Effect of different starting nematode densities on larval fitness

Nematode infections are highly costly to beetles. We observed significant treatment effects associated with different nematode densities on brood size, total brood weight, the number of eclosed adults and the fraction of larvae that went on to eclose as adults (Kruskal-Wallis, all p < 0.001, Table 1). In addition, we observed a significant negative relationship between the number of inoculated nematodes and mean brood size (r^2^ = 0.84, p = 0.03) and mean larval mass (r^2^ = 0.9, p = 0.02) (Figure 2). Brood size declined nearly 3-fold in broods with the highest nematode densities (Mean ± SE: 11.21 +/- 2.15 larvae/brood) compared to nematode-free broods (Mean ± SE: 27.25 +/- 3.03 larvae/brood), while mean larval mass declined roughly 15% from 0.181 +/- 0.006 g in broods without nematode infections to 0.156 +/- 0.007 g in broods where females were inoculated with 10,000 worms.

**Table 1:** Summary statistics for broods produced by worm-free females and females infected with different densities of nematodes prior to mating.

| *Infection level* | **Number of broods** | **Brood size (excluding failed broods)** | **Total brood mass** | **Mean larval mass** | **Number of eclosed adults** | **Fraction eclosed** |
| --- | --- | --- | --- | --- | --- | --- |
| **0** | 20 | 32.06 +/- 3.03 | 5.67 +/- 0.21 | 0.181 +/- 0.006 | 28.82 +/- 1.46 | 0.91 +/- 0.03 |
| **10** | 32 | 17.41 +/- 2.07 | 3.03 +/- 0.29 | 0.165 +/- 0.005 | 12.85 +/- 1.67 | 0.68 +/- 0.07 |
| **100** | 32 | 19.90 +/- 2.22 | 3.65 +/- 0.36 | 0.171 +/- 0.004 | 13.81 +/- 1.84 | 0.59 +/- 0.05 |
| **1000** | 32 | 13.41 +/- 1.88 | 2.62 +/- 0.28 | 0.162 +/- 0.005 | 5.88 +/- 1.54 | 0.36 +/- 0.07 |
| **10000** | 32 | 11.21 +/- 2.15 | 2.34 +/- 0.36 | 0.156 +/- 0.007 | 10.45 +/- 2.01 | 0.61 +/- 0.07 |
| **Kruskal-Wallis** |  | p < 0.001 | p < 0.001 | p = 0.09 | p < 0.001 | p < 0.001 |
| **r<sup>2</sup>** |  | 0.84 | 0.72 | 0.90 | 0.65 | 0.55 |
| **p value** |  | p = 0.03 | p = 0.07 | p = 0.02 | p = 0.10 | p = 0.15 |

**Figure 2:**
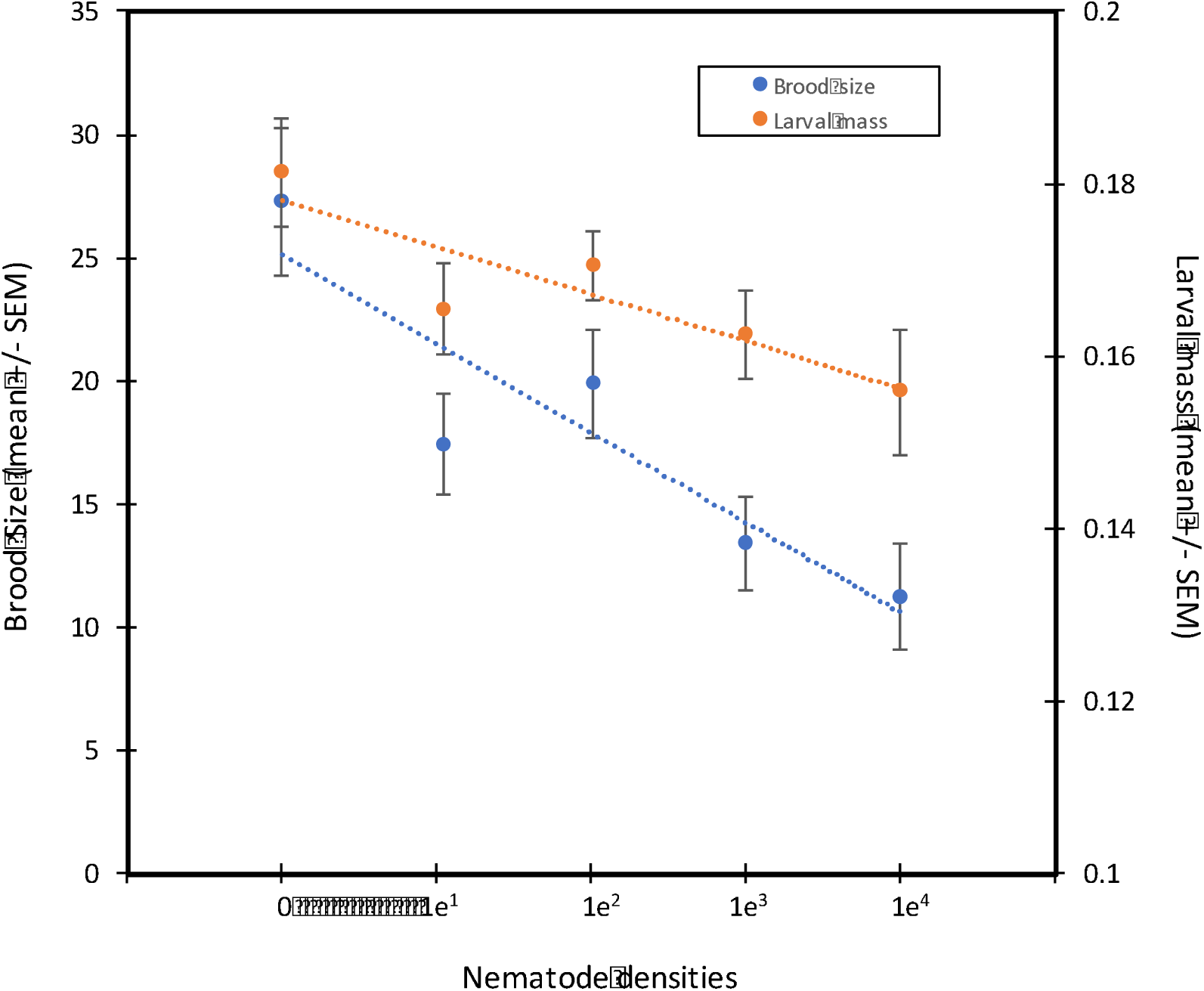
Decline in brood size and larval mass as a function of initial nematode numbers on mated females.

In addition to these direct negative effects of nematode infection, worms also altered the well-known trade off in *Nicrophorus* between brood size and average larval mass (Figure 3). Whereas there was a significant negative size:number trade-off in worm-free beetles (r^2^ = 0.50, p = 0.001), there was no such association in broods produced by females infected with worms at any of the treatment densities (all NS).

**Figure 3.**
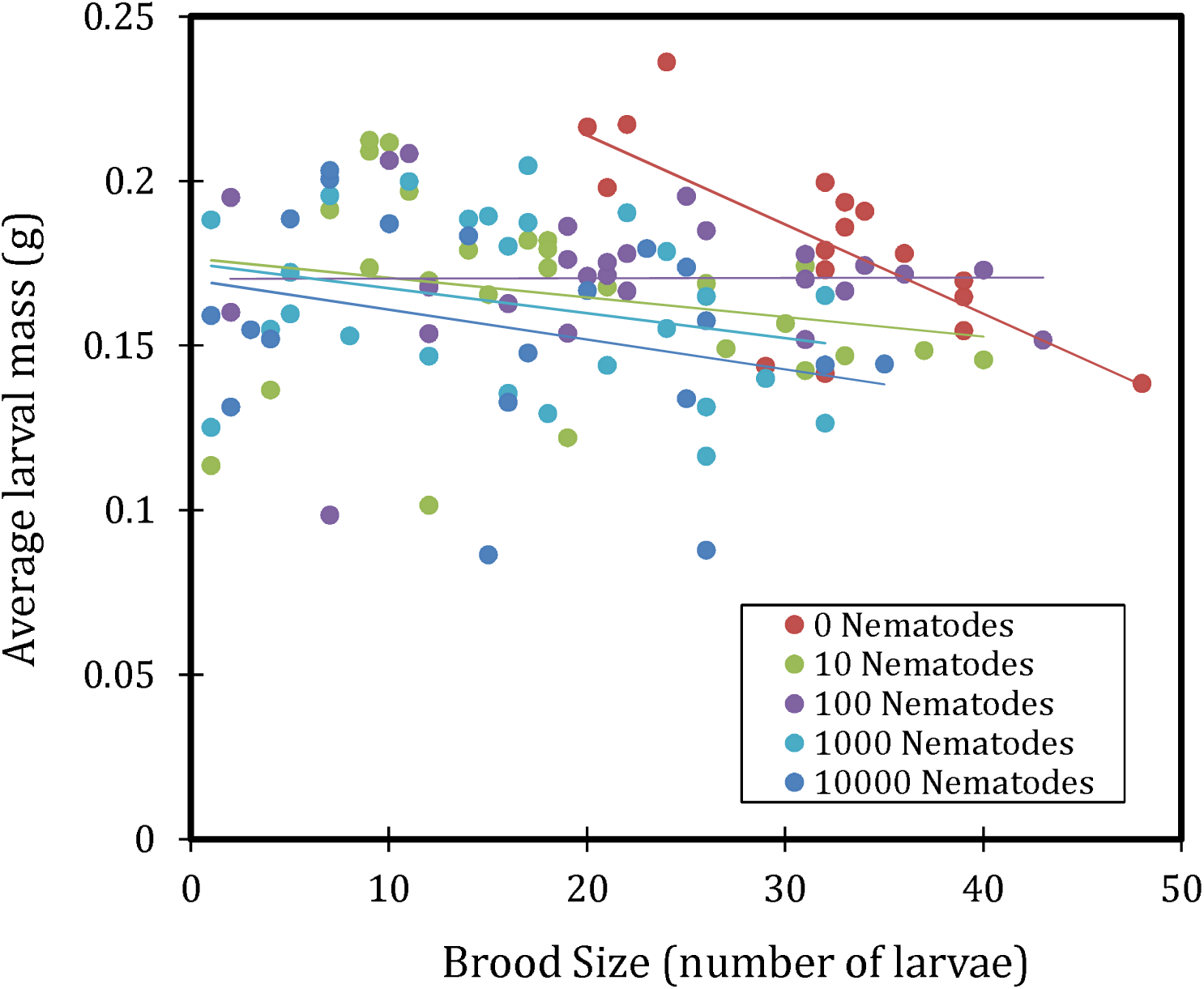
The relationship between average larval mass (g) and total brood size. Lines represent linear regressions. All are NS except for the nematode-free treatment.

### Transmission between sexes and from mothers to offspring

We inoculated male or female beetles (donors) with different nematode densities and then measured worm transmission to opposite sex recipients during mating. As shown in Figures 4A and 4B, we observed intersexual transmission of nematodes in both directions, although this varied with worm density and the sex of the donor. For both donor sexes, when the initial density of nematodes was 10, there was neither transfer nor retention of worms. Transmission occurred at all other initial worm densities.

**Figure 4.**
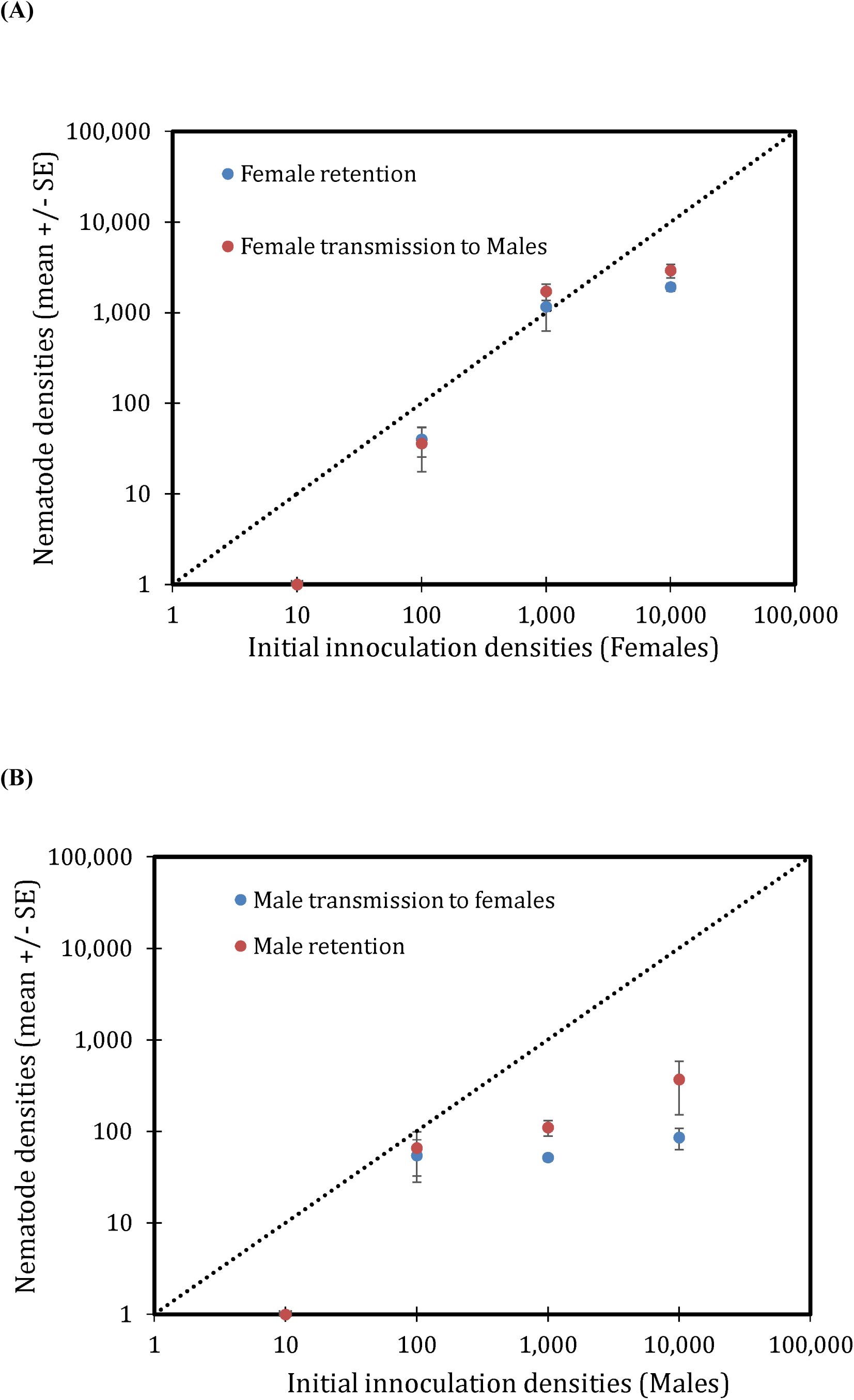
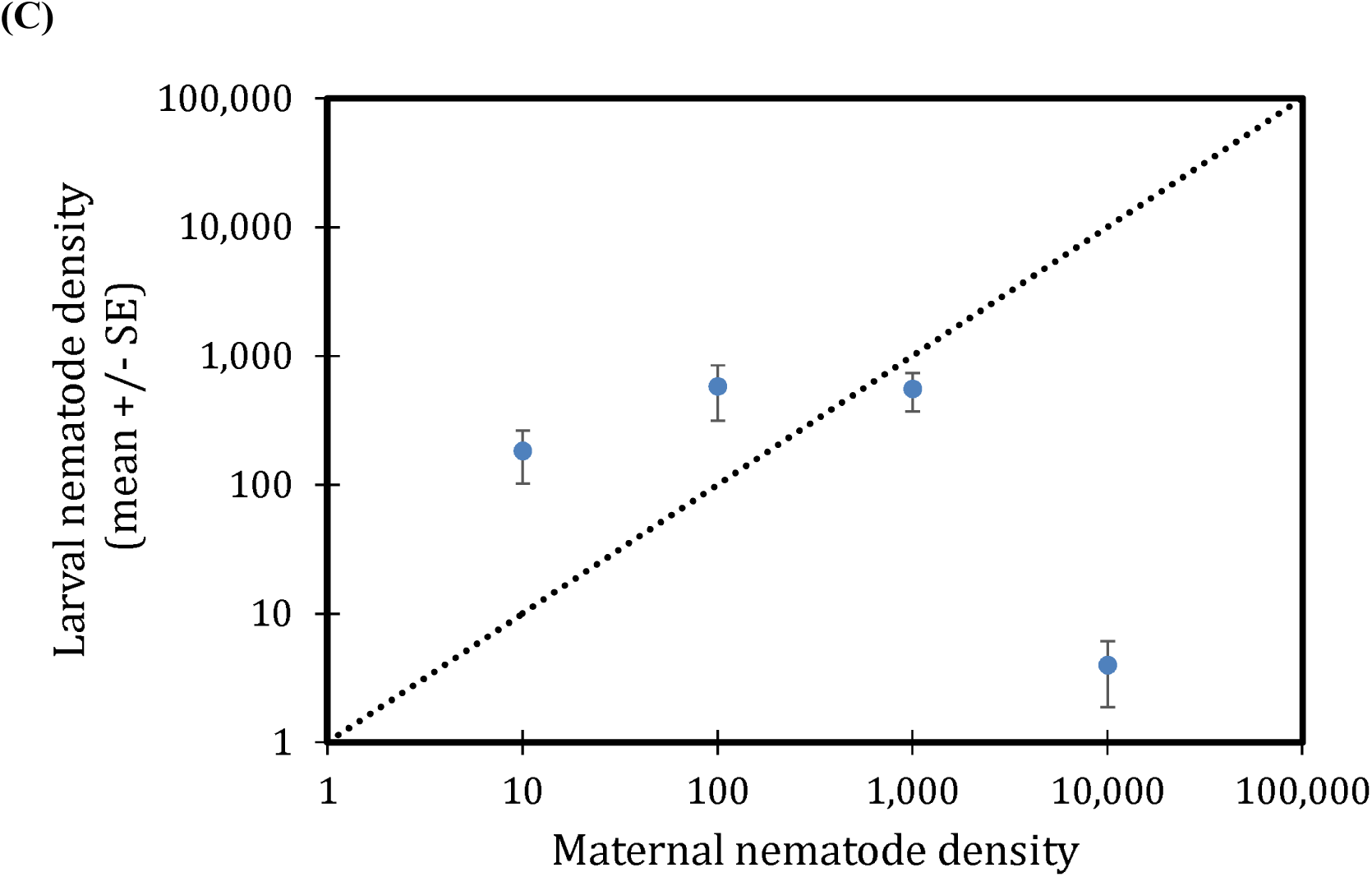
Nematode transmission between mated pairs and between generations. (A) transmission from females to males; (B) transmission from males to females; (C) transmission from mothers to offspring.

At inoculation densities of 100, worm numbers declined slightly in both male and female donors; however, transmission occurred effectively and there were no significant differences in the final worm densities of donors or recipients. Worm densities were retained at initial values when donor females were inoculated with 1,000 worms but were significantly reduced when the initial inoculum was 10,000 worms (t_5_ = -23.2, p < 0.001). Final worm densities in males and females (recipients and donors, respectively) did not differ at either inoculum density. In addition, there were no differences in final worm densities between beetles inoculated with 1,000 or 10,000 worms, suggesting an apparent carrying capacity of roughly 2,000 worms per adult beetle (mean +/- SE: 1770 +/- 238.3).

For male donors, both transmission and retention were reduced compared to female donors. At initial densities of 1,000 or 10,000, we observed significant reductions in donor and recipient worm densities overall (1;000: t_5_ = -56.64, p < 0.001; 10,000: t_5_ = - 83.5, p < 0.001). There were no differences in final worm densities at all inoculum sizes >10, suggesting a carrying capacity of approximately 125 worms/adult beetle (mean +/- SE: 123 +/- 41.6). Although both sexes can transfer nematodes to the opposite sex during mating, female transfer and retention is approximately 10X higher in females than males (t_34_ = 6.3, p < 0.001).

Next, to measure transmission from mothers to offspring, we inoculated mated females with different densities of worms and then allowed them to rear broods, after which we quantified the number of nematodes on eclosing pupae (Figure 4C). For all inoculum sizes other than 10,000, we observed significant transmission from mothers to larvae (one-sample t-test: 10: t_6_= 2.78, p = 0.039; 100: t_10_ = 2.43, p = .036; 1,000: t_13_ = 3.02, p =0.01). In addition, worm densities on eclosing larvae were significantly or marginally greater than maternal inoculum densities (one-sample t-test: 10: t_6_= 2.6, p = 0.047; 100: t_10_ = 2.0, p = .072; 1,000: t_13_ = -2.394, p = 0.032). Finally, we observed no significant differences in the final densities of worms on eclosing larvae from the 10, 100, and 1,000 treatments (one-way ANOVA: F_2,30_ = 0.79, p = 0.463), with a mean of approximately 500 nematodes per eclosed individual (mean +/- SE: 494.1 +/- 120.2). Unexpectedly, we found negligible transmission when mothers were initial inoculated with 10,000 nematodes, a likely artifact attributed to the extremely high rate of larval mortality in this treatment group (brood success ∼ 7%).

## Discussion

Our results provide the first evidence for an association between *Nicrophorus* beetles and the nematode *R. regina* or a novel species closely related to *R. regina,* a species only previously known from the haemocoel of scarab beetles (38). Because it causes high mortality in scarabs and releases bacteria when it infects these beetles, *R. regina* has been characterized as an entomopathogen that feeds on the bacteria that proliferate within the beetle cadaver (39, 40). Here, although we find that *R. regina* harms *Nicrophorus*, its behavior and transmission is more consistent with phoresy. In particular, we observed massive population growth of nematodes on the carcass itself (possibly due to consumption of bacteria on the carcass) and also conspicuous worm nictation upon disturbance, a behavior commonly seen in phoretic species (41). Our field collections reveal that this species is maintained in high, although variable, densities in male and female wild beetles (Figure 1) while further studies (YW unpublished) have shown that they are also stably maintained within laboratory populations of burying beetles at even higher densities. Although *Nicrophorus* nematodes are believed to have no or marginal effects on beetle fitness, our results indicate that this is not the case. Worm infections cause significant harm to beetles and the extent of this harm scales with worm density for both brood size and mean larval mass (Figures 2 and 3, Table 1), two central measures of adult and larval fitness, respectively. In addition, nematodes are transferred at high rates between adults and from parents to offspring (Figure 4).

We find strong density-dependent effects of nematodes on *N. vespilloides*. However, even though we see a significant scaling between worm numbers and e.g. brood size and mean larval mass, much of the maximum cost observed at the highest worm density (10,000) is already observed at the lowest inoculum size we used (10 worms). In other words, of the ∼ 50% decline in brood size in beetles inoculated with 10,000 worms, some 80% of this decline is already apparent in beetles inoculated with only 10 worms. This result is consistent with the idea that worms are proliferating extensively on the carcass where they can then go on to infect larvae, which seems to occur whether the initial number of colonizing worms is high or low. This result also explains why nematode transmission from parents to offspring has no lower threshold (Figure 4C), in contrast to the threshold of ∼100 worms needed for transmission between breeding adults (Figures 4A and 4B). Equally, while the densities of worms on larvae tends to exceed the inoculum density on females (consistent with proliferation), this is not the case for intersexual transmission, suggesting that nematode reproduction does not occur in this context and that worm transmission between adults suffers from stochastic loss if worms are initially rare.

Less clear are the factors that are responsible for the harm worms cause to beetle larvae. Nematodes can potentially cause indirect harm to beetle larvae by competing with them for space or resources, or possibly, by physically interfering with larvae while they consume the carcass. Phoretic nematodes are bacterivores, so direct resource competition with beetle larvae seems unlikely, unless some part of beetle nutrition is also microbial (directly or indirectly) (42). Competition for physical space may occur if nematode densities are sufficiently high to prevent larval feeding or access to parts of the carcass. This type of interference could also explain the absence of a trade-off between brood size and larval mass (Figure 3), since much of this effect is driven by the reduction of larval size in smaller broods (< ∼10 larvae/brood). Direct harm could possibly arise at different stages of development. Eggs could be pierced, something observed by *Nicrophorus* phoretic mites, *Poecilochirus carabi* (43), or otherwise damaged by nematodes. Larvae could also be directly harmed by worms during their growth (44). It is notable that worms are not only transported on the surface of beetles, but are also recovered from within the digestive and reproductive tracts (Figure 1), indicating an ability to invade host tissue (45, 46). Internalized worms may obtain nutrients from the larvae or otherwise hinder their growth and development (45). At present, this remains unknown because we lack intermediate samples of larvae themselves, and instead have focused on characterizing the transmission route of worms from mature adults through to newly eclosing adults. It will be interesting in future work to sample worm densities in developing larvae to better understand how and when worms inflict their damage.

Although our results make clear that *R. regina* is common in field-caught beetles and can persist through a complete beetle life-cycle, there are important limitations to our study. Most notably, our fitness experiments were carried out in the lab in the absence of other species that could either mitigate or exacerbate the harm caused by nematodes. *Nicrophorus* beetles carry other phoretic species: many different species of mites (31) and possibly other species of nematodes (38, 47). While there is no evidence of simultaneous carriage of different nematode species, this has been observed in other insect:nematode associations and is possible here, too (38). On the other hand, mites and nematodes always co-occur in *Nicrophorus*. Wilson and Knollenberg (1987) found that the effects of mites varied from harmful at high densities to neutral or even beneficial at lower densities. They also found that mites reduce the burden of nematodes on eclosing adults by up to 6-fold, from ∼ 18,000/individual to ∼ 3,000/individual (30). This reduction could have different causes, from direct consumption of nematodes to other types of interference competition; regardless, their experiments make clear the importance of examining the effects of phoretic species in the context of the entire community. This includes mites and nematodes, but also should include the microbial species that live on and within the beetles and carcass, and also the microbes that are carried within the nematodes, especially because these bacteria may be directly associated with nematode entomopathogenicity (44, 48)

Phoresy is extremely common in nematodes (49). Our results show that the phoretic worms of *N. vespilloides* are detrimental to beetles even though they rely on beetles for transmission to new resources. However, this harm may be inevitable and unavoidable if worm transmission requires proliferation. It will be interesting to determine if this is similarly true for other phoretic associates of animals.

## Authors’ contributions

YW and DR conceived the ideas and designed methodology; YW collected the data; YW and DR analysed the data and co-wrote the manuscript. Both authors contributed critically to the drafts and gave final approval for publication.

